# Genome-Wide Identification, Characterization and Expression Analysis of Non-Arginine Aspartate Receptor like kinase gene family under *Colletotrichum truncatum* stress conditions in Hot pepper

**DOI:** 10.1101/2020.01.23.916528

**Authors:** R Srideepthi, MSR Krishna, P Suneetha, R Sai Krishna, S Karthikeyan

**Affiliations:** Department of Biotechnology, Koneru Lakshmaiah Education Foundation, Guntur, Andhra Pradesh, India; Institute of biotechnology, Professor Jaya Shankar Telangana State Agricultural University, Hyderabad, India; Crop production, ICAR-National Rice Research Institute, Cuttack, Odisha, India

**Keywords:** Pattern Recognition Receptors (PRRs), Autophosphorylation, Downstream Signaling, *Colletotrichum truncatum*, defense responses.

## Abstract

Receptor Like kinases (RLKs) are conserved upstream signaling molecules that regulate several biological processes, including plant development and stress adaptation. Non arginine aspartate (non-RD) an important class of RLKs plays a vital role in disease resistance and apoptosis in plants. In present investigation, a comprehensive Insilco analysis for non-RD Kinase gene family including identification, sequence similarity, phylogeny, chromosomal localization, gene structures, gene duplication analysis, promoter analysis and transcript expression profiles were elucidated. In this study twenty six genes were observed on nine out of twelve chromosomes. All these genes were clustered into seven subfamilies under large monophyletic group termed as Interleukin-1 Receptor-Associated Kinase (IRAK) family. Structural diversity in genomic structure among non-RD kinase gene family were identified and presence of pathogen induced *cis* regulatory elements like STRE, MYC, MYB,W box were found. Expression profiles of genes involved in providing resistance to anthracnose pathogen *Colletotrichum truncatum* in hot pepper were analyzed at different infective stages in both resistant and susceptible genotypes. Among twenty six genes, *CaRLK1* gene belonging to LRRXII subfamily was up regulated under severe stress after infection in resistant genotype PBC-80. This integrative approach has helped us to identify candidate genes involved in disease resistance which would be helpful in future crop improvement programs.

## Introduction

One of the major challenges in the 21^st^ century, to global food security and agricultural sustainability is to develop economically important high yielding varieties which are stable with broad-spectrum of resistance. Hot pepper is a commercially important vegetable crop grown worldwide for its indispensable nutritional and therapeutic values, but year by year its production has been reduced due to several biotic stresses [1]. Reason might be due to existence of limited resistant cultivars, existence of variability within the *Colletotrichum truncatum* species with erratic pathogenic ability with respect to different hosts, climatic conditions and commercial pesticides used remained as an unsatisfactory measures for its effective control [2]. As a part of its development of anthracnose resistant hot pepper remained as one of the major tasks to be resolved by modern agriculture practices prevailing worldwide. Congruently knowledge of understanding the defensive signaling mechanisms employed by resistant plants while encountering the attacking pathogen by advanced molecular and computational techniques paved a ray of hope to address these challenges [3].

RLKs are surface localized receptor-like protein kinases employed by Pattern recognition receptor (PRRs) proteins of plants innate immune system as a primary defensive response [4]. These are involved in signal perception from pathogen by ectodomain, then transduction of signal by transmembrane region and then activation / deactivation of signaling cascade by kinase domain. Classification of ecto-domain was done based on the type of ligand binding specificities [5] and endo domain based on presence or absence of conserved arginine residues present immediately preceding to aspartate in catalytic domain VI of kinase domain [6]. Among various types of RLKs, predominantly non-RD RLKs were allied with innate immune receptors that recognize conserved microbial signatures and activate pattern triggered immunity (PTI) involved in disease resistance [7]. Till now 35 genes in Arabidopsis and 328 genes in Rice were identified to possess non-RD class of kinase receptor proteins. Functionally characterized non-RD kinases like, XA21 BSR1 and XA26 from Rice and FLS2, EFR from Arabidopsis thaliana were known to be effective against bacteria [8] while Pi-D2 gene of lectin (non-RD) kinase was found to express broad-spectrum resistance against *Magnaporthe grisea* [9]. *LecRK-VI*.*2* from *Arabidopsis thaliana* [10] and WRKY from tomato [11] serve as a potential link in providing resistance against bacterial and fungal pathogens. Hence due to limited availability of literature on non-RD kinases in hot pepper and available free online software motivated us to identify the presence of candidate non-RD genes associated with disease resistance through comparative genomics.

## Materials and methods

### Identification of transmembrane receptor kinases in hot pepper

Identified and known RLKs from sequenced plant genomes like Arabidopsis, Rice and tomato were collected as query sequences from NCBI (https://www.ncbi.nlm.nih.gov). Collected query sequences were submitted to blast search against hot pepper CM-334 variety genome datasets (http://www.solgenomics.net) version 1.55 of *capsicum annuum*. Protein sequences sharing more than 50% homology with query were selected for further analysis. The presence of signal peptide was determined by SignalP 4.0 Server (http://www.cbs.dtu.dk/services/SignalP/)[12]. Extracellular ligand-binding domain, transmembrane, and intracellular kinase domain were filtered by online domain search databases SMART (http://smart.emblheidelberg.de/) [13] and Pfam with inbuilt HMMER search platform [14]. While Conserved motifs were identified using the MEME program (http://meme-suite.org/index.html). This program was run with default settings except to set a maximum ten in number with the three hundred widths of motifs [15]. Based on the conserved motifs and phylogenetic analysis, receptors like kinases in hot pepper were bundled.

### Phylogenetic analysis of transmembrane receptor kinases

All known and recognized transmembrane receptor protein kinases (RLKs) belonging to model plant species like *Solanum Lycopersicum* [16], *Arabidopsis thaliana* and *Oryza sativa* [17] were taken together for phylogenetic analysis. Identified and characterized reference proteins at least one from each family and subfamilies belonging to respective clades of three model plants along with identified putative gene CaRLK Fasta sequences were submitted to MEGA software version 6.0 [18]. All sequences of CaRLK genes were submitted to multiple sequences alignment using Multiple Sequence Comparison by Log-Expectation (MUSCLE) with default values. Then phylogenetic tree was constructed using the Neighbor-joining (NJ) method with thousand numbers of bootstrap replications.

### Analysis of CaRLK Physiochemical Properties

Physiochemical parameters of putative CaRLK gene including molecular weight, isoelectric point, number of amino acids, aliphatic index, and grand average of hydropathicity (GRAVY) score was determined using online ExPASy programs (http://www.expasy.org/). Subcellular locations of CaRLK proteins were predicted using the online Plant-PLoc tool (http://www.csbio.sjtu.edu.cn/bioinf/plant-multi/) program.

### Chromosomal Distribution and Duplication of CaRLK Genes

Information regarding position and location of chromosomes on which CaRLK gene members were present was derived from hot Pepper Genome Platform (PGP) (http://Hotpeppergenome.snu.ac.kr/). Genes were mapped onto chromosome at their genomic position and drawn manually. Duplicated genes were identified by Blast P search against each other when both their identity and query coverage was >80% of their partner sequence [19]. Tandem duplication in genes was identified by occurrence of homologous genes located in single region(<100kb) within a chromosome, while segmental duplication occurs among homologous or non-homologous genes with > 1kb in length and more than 90% sequence similarity dispersed but present on same or different chromosomes from the same clade as described by Feng et al. (2017) [20]. Consequently, non-synonymous (Ka) and synonymous substitution (Ks) among duplicated CaRLK gene pairs were calculated using PAL2NAL (http://www.bork.embl.de/pal2nal/). The divergence time of the duplicated gene pairs was calculated as described by Koch et al. (2000) [21].

### Gene structure and Cis-regulatory element analysis

Gene structure was elucidated based on the relationship of the coding sequence and its corresponding genomic DNA sequence by GSDS 2.0 (http://gsds.cbi.pku.edu.cn/).Cis-acting regulatory elements of genomic DNA sequences of 3000 bp 5’ upstream region was mined from the Sol Genomics Network database. Promoter sequences obtained were submitted in Plant Care Database http://bioinformatics.psb.ugent.be/webtools/plantcare/html/) individually. Conserved biotic-stress responsive elements in hot pepper were predicted, as described by Diao et al. (2018) [22].

### Primer Design

Genomic and its Coding sequences (CDS) of deduced RLK hot pepper proteins were retrieved from Sol Genomics Network (https://solgenomics.net/). Primer sets for RT-qPCR were designed in 3 and 5 untranslated regions of individual genes to avoid non-specific amplification using Prime Quest Tool (http://eu.idtdna.com/PrimerQuest/Home/Index) [23]. For all primers Ubiquitin 3 was used as reference gene. Genes with accession numbers and code assigned in the study (S1-Table).

### Plant Material, Fungal Strains and Stress Treatments

Seeds of two hot pepper genotypes pbc-80 (anthracnose resistant) from National Bureau of Plant and Genetic Resources (NBPGR), Hyderabad and Pusa Jwala (anthracnose susceptible) from Horticulture Research Station, Lam Farm, Guntur, Andhra Pradesh were collected for the study. Seeds were scattered in black trays containing autoclaved blend of peat and vermiculite (2:1 v/v) along with micronutrients mixture. Seedlings were raised and watered regularly in a greenhouse under controlled conditions i.e., 16 h light/8 h dark photoperiod at 27 °C throughout the day and 21°C during the night. Fungi were isolated from fruit rot infected hot pepper samples by single spore isolation technique. Isolate *Colletotrichum truncatum* was cultured on Oatmeal agar medium with pH 7.0 at 25±2°C. Spore suspension was prepared from seven days old culture and sprayed on seedlings by artificial inoculation method as described by Mishra et al (2017) [24].with 5x 10^5^spores/ml concentration. Three-week old hot pepper seedlings were taken for experimental studies. Genotypes PBC-80 and Pusa Jwala sprayed with conidial suspension were considered to be treated and those trays sprayed with autoclaved distilled water as control. Each treatment was maintained with three replications.

### RNA Extraction

RNA was extracted from leaf tissue of stressed and control genotypes according to acid guanidinium thiocyanate-phenol-chloroform extraction method [25]. Leaf tissue (100 mg) was ground into fine powder by using liquid nitrogen in mortar and pestle. Fine powder was transferred carefully into 2 ml eppendorf tubes with extraction buffer and centrifuged at 12,000 rpm at 4° C for 10 minutes. Supernatant was collected and equal aliquots of chloroform was added to it and centrifuged for 10 minutes at 12,000 rpm. To upper aqueous layer ice cold propanol and 1.5 M NaCl was added and incubated at 4 ° C for 5 min. Then tubes with solution was centrifuged for 10 minutes at 12,000 rpm and supernatant was discarded. The resultant pellet was subjected to ethanol wash and allowed for air drying. Pellet was dissolved in DEPC water. RNA samples were treated with DNase I (RNase-free) (Takara-cat # 2270B) to remove residual genomic DNA. The purity and integrity of RNA were determined by calculating A260/A230 and A260/280 absorbance ratio. RNA was visualized in 1.5 percent agarose gel after electrophoresis.

### Quantitative Real-Time PCR

cDNA was produced in triplicates from three μg of RNA by prime script 1st strand cDNA synthesis kit (Takara-cat # 6110A). Gene expression was scrutinized in reverse transcription-quantitative polymerase chain reaction (RT-qPCR) (QuantStudio3-Applied Biosystems). Twelve μl reaction mixture (Sybr Green—6 μl, cDNA—2 μl, Forward Primer—1 μl, Reverse Primer—1 μl, DEPC water—2 μl) was loaded in 96-Well Reaction Plates enclosed with microamp Optical Adhesive Film (Applied Biosystems). Standard cycling parameters and baseline thresholds were set manually along with UBI-3 as a reference gene. CT values were calculated by using software. Proficiency of RT-qPCR reaction for every single RLK gene was deliberated from standard curve gained by serial dilutions of pooled cDNA. Relative gene expression among the samples was calculated using the 2-DDCT method [26]. In this method, an estimated level of gene expression was purely based on a hypothesis of 100% PCR efficiency of reference and target genes. RT-qPCR products were visualized on 2% agarose gel. The presence of distinct single band with expected amplicon size was considered as specific amplification [27].

### Statistical Analysis

Expression studies for all the four samples (pbc-80 control, pbc-80 stressed, pusa jwala control and pusa jwala stressed) were carried out at three biological replicates, two technical replicates and were retained at each point of time intervals. Data were represented as mean ± standard deviation. All the data were tested for significance between resistant and susceptible genotypes using analysis of variance (ANOVA) at 5% probability. Relative gene expression was generated using heat mapper (http://heatmapper.ca/).

## Results

### Identification and annotation of transmembrane receptor kinases

Blast P homology searches revealed 3,150 hits with pepper genome database. Among them 1,208 hot pepper sequences were found to share more than 50 percent homology with a sequence length ranging from 100-2000 amino acid [28]. From PDB database optimized length of known receptor like kinase sequences ranged from 100-1000 amino acids [S10]. Hence non redundant sequences within the optimized range were considered for further study while remaining redundant sequences were removed manually. A Total of 164 Sequences with typical domain organization viz., presence of signal peptide, varying extracellular domains, a transmembrane region and an intracellular kinase domain were retrieved from respective databases. Based on Conserved motifs type of ectodomain was distinguished and details were illustrated in (S6 Fig). Naming to each gene was specified with the first two letters indicating *Capsicum annuum* (Ca) third letter representing the non-RD class of transmembrane receptor-like kinase (RLK) family followed by serial numbers [29].

### Phylogeny Analysis

Phylogenetic analysis revealed the existence of seven gene families under respective clades in which each clade contains one gene from tomato, and Arabidopsis (Fig 1). Phylogeny grouping revealed the evolutionary relationship of each gene family and their respective subfamilies under respective clades. Pictorial illustration revealed the occurrence of 1. Cysteine-rich receptor-like kinases (Stress antifung) represented in pale green color. 2. Wall Associated Kinases (LRK10L2, LRK10L1) in red color. 3. L-type Lectin receptor like kinases (lectin Leg B) in violet color. 4. G-type lectin receptor like kinases (SD1a, SD2b, SD3) in green 5. Lys M receptor like kinases in pink color. 6. Malectin receptor like kinases in chocolate brown color. 7. Leucine-rich repeat (LRR) receptor like kinases – LRR Ia, LRR-II, LRR-III, LRR-IV, LRR-V, LRR VIII, LRR IX, LRRXI, LRRXII in dark green color were found. In hot pepper extracellular domains containing RLKs similar to C-type lectin (sky blue), URK1, PERK, extensin (yellow) receptor like kinases were not observed in our analysis.

**Fig. 1.**
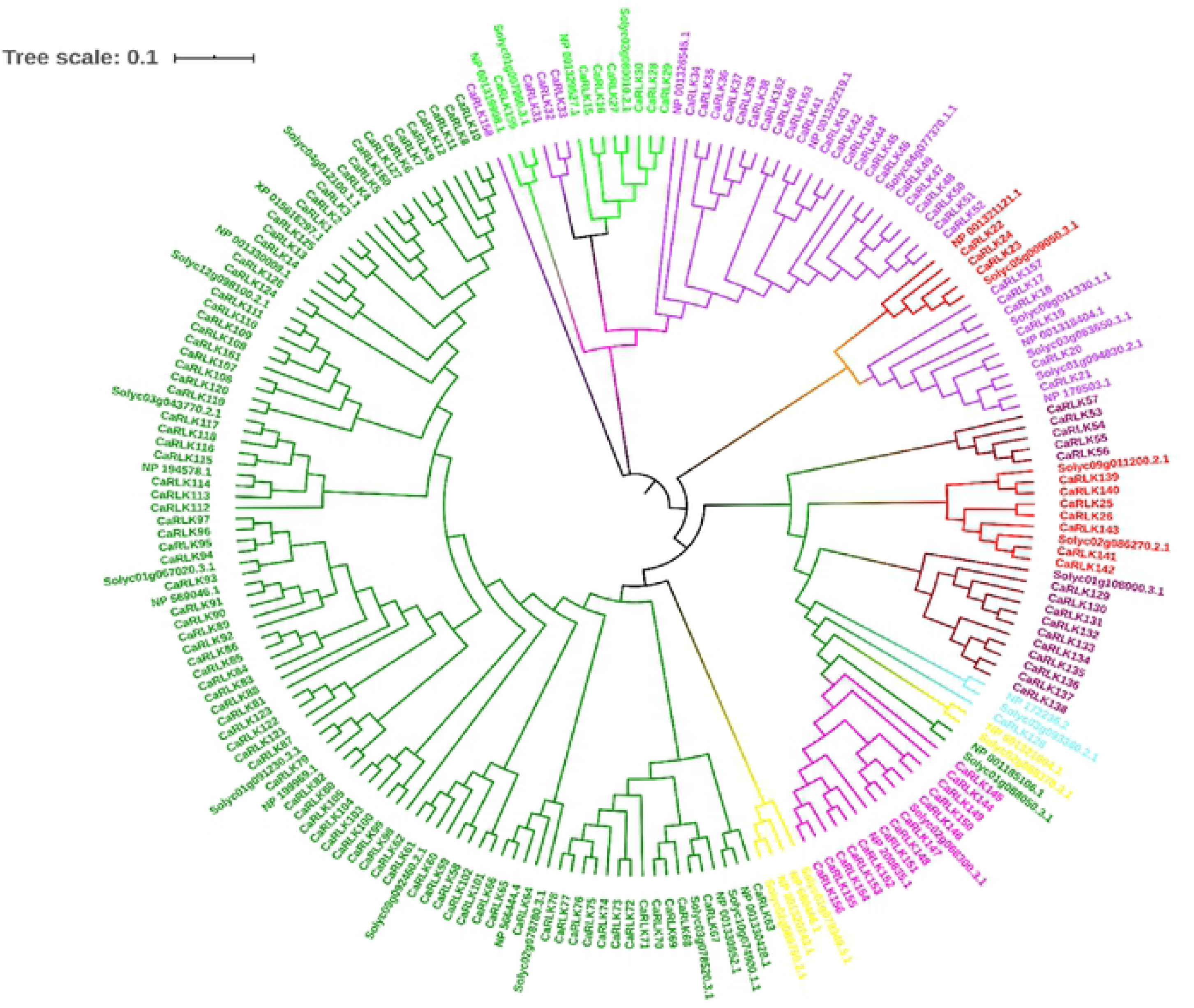
Phylogenetic analysis of Receptor like kinase gene family from hot pepper. The phylogenetic tree was constructed using neighbor-joining (NJ) method by MEGA6.0.Subfamilies was specified in different colors

### Classification of Kinase Domains

Based on kinase domain organization CaRLKs were classified into three types. Based on the presence or absence of the conserved residue Arginine (R) present adjacent to Aspartic acid (D) in subdomain VI, conserved lysine (K) residue and Aspartic acid (D) in subdomain II and VII kinase regions. Presence of Arginine (R) adjacent to aspartic acid as RD Kinases those lacking R as Non-RD Kinases and lack of any one or more of the K/D/D residues were classified as RD minus. Finally, 26 protein sequences were found to possess Leucine, cysteine, phenylalanine, Glycine, and Serine residues substituted in place of arginine. In Supplementary Table (S2) distribution of conserved residues in II, VI and VII domains of 26 genes with respect to their corresponding ectodomain and respective clades fitting to them were given in detail. Multiple sequence alignment of twenty-six genes (S5 Fig) were predicted to share a common linkage as they are descended from a common ancestor. However all 26 CaRLK genes were predominantly grouped under respective LRRXII, LRRXI, SD1a, SD2b, SD3, LRK10L2 and WAK/LRK10L1 clades in their phylogenetic relationship.

### Physiochemical Properties

Non-RD receptor-like protein kinases vary significantly with respect to their structural and physical properties. Genes CaRLK6 with the highest number (1789) of amino acids and CaRLK23 with the lowest number of amino acids (620) was observed while CaRLK17 with high molecular weight (954.89) and CaRLK 23 with low molecular weight (70.11) respectively. Consequently, a wide variation was observed in their isoelectric point (PI) ranging from 5.69 (CaRLK14) to 8.63 (CaRLK10). Instability index ranged from 28.84 - 46.15 where 19 genes were considered to be stable as they exhibit instability index value less than 40, whereas the rest 7 among 26 non-RD RLKs were considered as unstable. Grand average hydropathicity (Gravy) values ranged from -0.345 (CaRLK19) to +0.12 (CaRLK10) inferring the presence of both hydrophilic and hydrophobic amino acids. Aliphatic index determines the thermal stability of a protein. Among 26 proteins high aliphatic index was observed (CaRLK 19-79.07 to CaRLK 7-111.49) indicating that all are thermally stable with more number of hydrophobic amino acids in their structure. Most of them were found to be localized in extracellular spaces of cell membrane followed by chloroplast, nucleus, and mitochondria. Physiochemical Characteristics of each protein were given (S3 Table) in detail.

### Chromosomal Localization

The inherent ability of a host to defend against pathogens mostly depends on the occurrence of resistance genes and activation of their signaling mechanism present on chromosomes. A total of twenty-six putative genes belonging to non-RD class were aligned on 9 out of 12 chromosomes. Chromosomes 3, 9 and 10 were devoid of non-RD genes, while in contrast a maximum number of seven genes were observed on chromosome 2. Thirteen genes - CaRLK 9,11, 12, 13 on second, CaRLK6, 7 on fourth, CaRLK 2, 8, 10 on sixth while CaRLK3, 4, 5 on fifth and CaRLK 1 on eleventh chromosome belongs to LRRXII subfamily respectively as shown (S4). Single non-RD genes CaRLK 19, 20 and 24 were aligned separately on each of Chromosome 1, 8 and 12 were clustered under SD3, SD2b, and LRK10L2 subfamilies were aligned under same clade respectively. Four genes CaRLK 15, 17, 18, and 21 from same G-type Lectin subfamily located on chromosome 7 were scattered on different clade branches as shown phylogeny figure.

### Duplication analysis of CaRLK genes in hot pepper

The evolution of gene duplication reveals results in family expansion and the occurrence of novel genes in a genome. Three pairs of genes showed tandem duplication CaRLK 27/28 and CaRLK 13/10, 14/15 while only one paralog pair CaRLK17/18from CRLK was identified to exhibit segmental duplication (S5). All three gene pairs of tandem duplication type possessed Ka/Ks ratios less than 0.5 indicating that genes experienced purifying selection pressure. While gene pair CaRLK 17&18 showed Ka/Ks ratios < 1.0 implying positive mode of selection. The divergence time of non-RD genes exposes the duplication events started from 5.86 Mya and continued up to 3.77 Mya in evolution.

### Gene structure analysis

Gene length varied from 2,224 bp (CaRLK 24) to 21,556 bp (CaRLK 6). Moreover, genes with either positive or negative sense strand as a template to the coding regions are depicted in Supplementary Table (S3). Various exon-intron positions were compared to gain insight into possible mechanisms of structural diversity existing among Non-RD kinases in *Capsicum annuum.* In this study introns varied from 0-12 in number. CaRLK 6 gene from LRRXII family showed a maximum of 12 introns. While in contrast a total of four genes (CaRLK 17, 18, 20 and 21) from G- type lectin and single gene CaRLK 13 from LRR type family were found to occur without introns in their structure. Genes from same family showed similar intron organization. Where Genes CaRLK 15, 16 from Stress-antifung subfamily showed six number of introns. While contrarily genes from LRR family possessed varied introns like ten genes (CaRLK 1, 2, 3, 4, 5, 8, 9, 14, 19 and 23) with single intron. Four Genes (CaRLK 10, 11, 22 and 24) with two introns and three genes (CaRLK 7, 25, 26) with three introns were observed in their structural organization. Moreover LRK10L2 clade members CaRLK23 with single intron and CaRLK 22, 24 with two introns also showed varied introns organization in their structure

### Cis-regulatory element analysis

*Cis-*regulatory element analysis promotes a good insight to understand the expression patterns of a gene under various stress conditions, whose validation needs to be warranted. Major pathogen-induced *cis*-regulatory elements identified in hot pepper (S7). Among twenty-six non RD genes highest number of *cis*-regulatory elements (TGACG, STRE) known to be involved in defense and stress responses were found in CaRLK 3, 11 from LRR and CaRLK 16 from G-type lectin subfamily. Fungal elicitor and oxidative responsive *cis* regulatory elements viz., W-box, F-box, As1 and box4 were observed majorly in CaRLK 1, 7,8 from LRR and 22, 26 from WAK family

Correspondingly elicitor responsive element G-box and Abscisic acid signifying region ABRE were observed in abundance among promoter regions of CaRLK3, 15, 17, 23 and 26 genes. Whereas Myb and Myc binding sites responsible for triggering stress-responsive metabolic pathway were found to occur predominately in CaRLK21, CaRLK 1 and 15. Moreover, only 32 percent of genes showed the presence of TC-rich repeat and TCA element linked with salicylic acid and methyl jasmonic acid pathway. When compared to all genes CaRLK1 and CaRLK16 from LRR family were identified to hold more number of *cis*-regulatory elements and may associate intensely with defense responsive mechanism.

### Heat Map Analysis

Differential expression of three family genes at varied time interval was observed in resistant (pbc-80) and susceptible (PJ) Hot pepper genotypes (Fig 2).

**Fig. 2.**
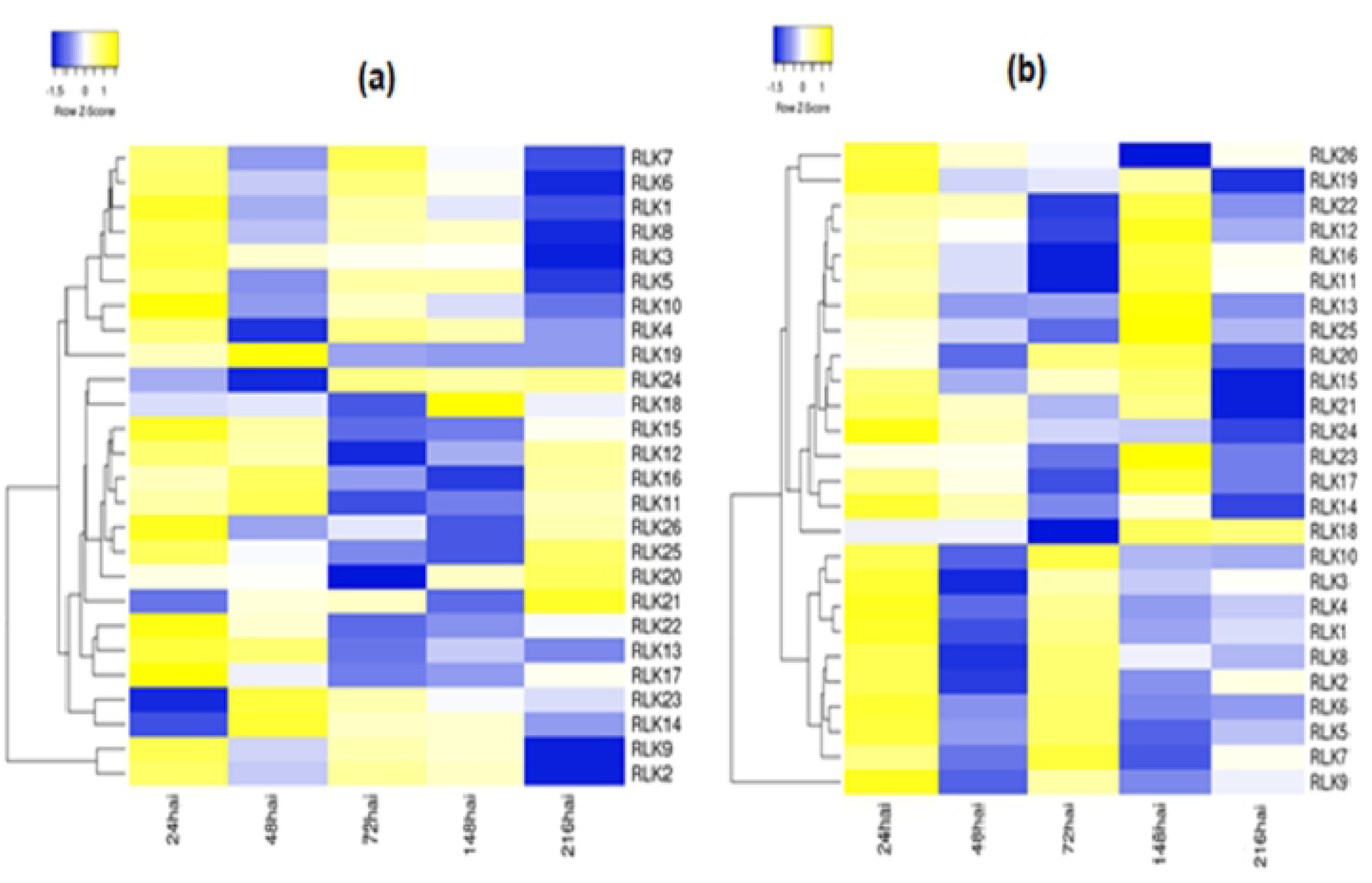
Differential expression analyses of CaRLK genes under biotic stress. Treatment in PBC-80 (a) and pusa jwala (b) Hot pepper seedlings. The color scale represents log2 expression values.

### Expression profiles of CaRLK genes at various stress treatments

At Initial stages of infection after 24 hours of inoculation, genes from LRR type (CaRLK 14, 1) G-type lectin (CaRLK15, 16 and 19) and Wall associated kinase (CaRLK 23, 24) showed significant up-regulation in Pbc-80 when compared to that of Pusa jwala cultivar (Fig 3.a). At biotrophic phase of infection, during colonization of subcuticular hyphae beneath the cuticle in resistant when compared to that of susceptible genotype after 48 hours of inoculation (Fig 3.b). Genes from LRR(14,6,3,1,2,9,10), G-type lectin (16,17,18,19) and WAK (23,24,26) families showed significant up-regulation while LRR (CaRLK2) exhibited down-regulation. During the necrotic phase, after 72 hours of inoculation defense-related genes (Fig 3.c). Gene CaRLK 2 from G-Lectin family, CaRLK 23, 25 from WAK, CaRLK 1, 9 and 14 from LRR families showed significant up-regulation. Furthermore genes CaRLK 17, 18 from G type lectin and CaRLK7 from LRR family have shown down regulation in resistant genotype when compared to susceptible genotype at a phase where extensive cell death of epidermal and mesophyll cells, instigating necrotic phase. In the present investigation number of genes belonging to LRR family was up-regulated in Pbc-80 indicating a potential defensive role. At formation of acervullus stage (216h after inoculation), genes from LRR (6,7,11,1,2,14), WAK (22,23,24,25,26) and G-Lectins (15,16,19,20) were significantly up-regulated While CaRLK 7,17 and 22 were down-regulated in pbc-80 when compared to pusajwala (Fig 3.d). Instability index of all genes up-regulated remained unstable in nature. At last stage, conidial dispersion after 216 h of inoculation. Genes from LRR family CaRLK 6, 1 and from LRRXII subfamily CaRLK8 showed a significant up-regulation. Whereas LRR (CaRLK 6) has down-regulated at severe stress conditions (Fig 3.e).

**Fig 3.**
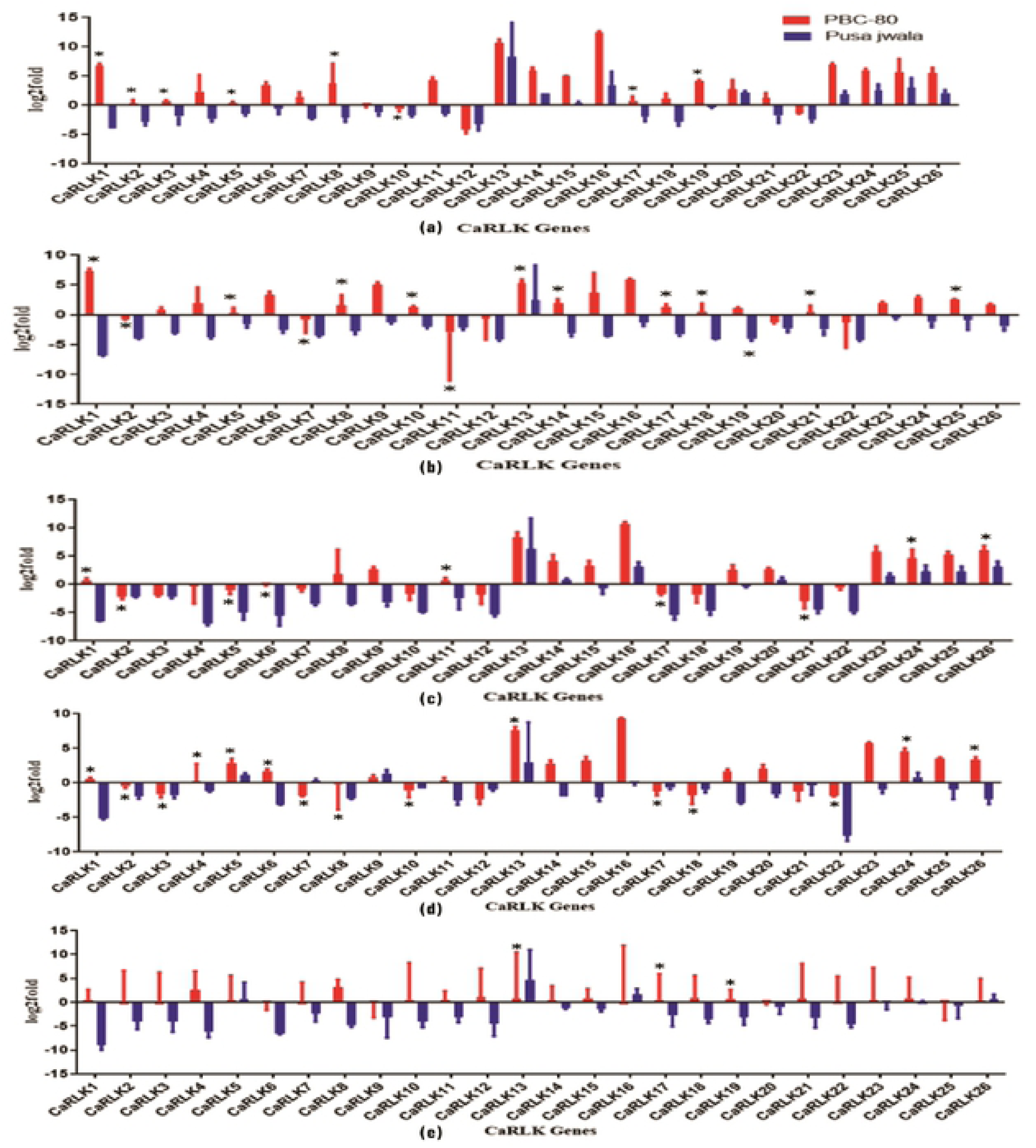
Expression profiles of CaRLK genes in leaf tissue in response to *Colletotrichum truncatum* at various time intervals (a) 24, (b) 48, (c) 72, (d) 148 (e) 216 hours after inoculation. Mean values and Standard Deviation for three replicates are shown.

## Discussion

Receptor-like kinases plays a vital role in plant development, signal transduction and defense responses [30]. Availability of sequenced genomes had facilitated the researchers to study various functional roles of RLK family genes in various stress adaptation procedures in many model plants like Rice, Arabidopsis, Tobacco, Wheat, Tomato, Soybean etc., [31]. Most of the candidate RLK genes involved in primary immune responses associated with disease resistance belong to non-RDclass kinases. Present investigation was aimed at genome-wide identification of non-RD kinases in hot pepper a vegetable crop with global agricultural and economic importance, whose production has been hindered by several biotic stresses [32]. A total of 8 and 35 percent of non-RD motifs were identified among IRAK family members in Arabidopsis and Rice [7]. This huge difference may be due to monocot-dicot diversification. Comparative phylogeny of hot pepper with model plants revealed the evolutionary existence of seven subfamilies from non RD class of RLK gene family. CaRLK genes were more closely related to genes from arabidopsis and tomato than rice, reflecting the fact that arabidopsis, tomato and pepper are eudicots and diverged more recently from a common ancestor [33].

Gene duplication events majorly include tandem, segmental and whole-genome duplications with substantial roles occurring in the evolution [34]. In hot pepper among 26 genes three pairs from chromosome 2 and 11 unveiled tandem duplications within LRR and WAK families evolved from common ancestor LRRXII family. In hot pepper, tandem duplication may signify LRRXII family lineage-specific expansion with novel gene expression. Whereas Hofberger et al. (2015) reported lineage-specific expansion of L-type lectin receptor kinase gene family by tandem duplication event in Brassicaceae. While only one pair from the stress-antifung subfamily of G-type lectin showed segmental duplication in the second chromosome. Segmental duplication majorly contributes to gene expression and plays an important role in immunity, growth and defense responses to external stimuli [20]. Results were in accordance with Cannon et al. (2004) [35] who reported a negative correlation between tandem and segmental duplication within Arabidopsis gene families. Furthermore, the substitution rate of non-synonymous (Ka) and synonymous (Ks) mutations was assessed to evaluate the selection pressures and divergence time succeeded among the duplicated CaRLK gene pairs (S4). LRR and WAK gene families formed by natural selection in hot pepper may hold extra functional members associated with speciation or adaptation. While single gene pair G-type lectin family showed positive selection and may be involved in functional diversification as described by Haung et al. (2016) [36].

Introns execute a major role in cellular and developmental processes via alternate splicing or gene expression regulation [37]. Five intron less genes were found in hot pepper, likewise, Yang et al. (2009) [38] also reported the presence of intron less genes in taxonomic species like *Arabidopsis thaliana, Populusdeltoides* and *Oryza sativa* representing their lineage-specific expansion with specific function in the evolution. In hot pepper among 26 non-RD genes only 10 genes had exhibited single intron in its structure. Xu et al. (2017) [34] also reported the presence of fifteen single intron genes in *Populus deltoides* with multiple functions. Genes present in a family usually contain the same structural organization but conversely, some genes from SD3, LRK10L2, and LRRXII subfamilies showed varied intron-exon organization in hot pepper. These variations may be due to substituted residues in conserved positions depicting the evolutionary changes occurring within a family [39].

*Cis-*regulatory elements are noncoding sequences that act as binding sites for transcription factors involved in proper spatiotemporal expression of genes containing them. In rice pathogen-induced, *cis*-regulatory elements like AS-1, G-box, GCC-box, and H-box were expressed and confirmed as markers for identification of resistance genes in response to fungal infection (Kong et al., 2018). While few genes CaRLK 17 and 3 from SD3 and LRRXII subfamily were found to possess those genes and may also imply the same function. The presence of *Cis*-regulatory elements like F/Sbox, W box, TGACG and MYB binding site in promoter regions are involved in the stress-inducible defense gene in Maize [40].

In this investigation, we had analyzed differential expression patterns of particular gene during various temporal stages based on disease progression studies (Fig 4). Infection stages of *Colletotrichum truncatum* like spore adhesion, germination, and penetration by appressorium (24h), subcuticular colonization of hyphae (48h), aggregation of mycelium (72h), Acervullus formation (148h), Conidial dispersion (216h) after inoculation on leaf surface of hot pepper seedlings was observed. Gene CaRLK 1 showed the highest expression in response to *Colletotrichum truncatum* even after 216hours of infection. Our results were in accordance with Sakamoto et al. (2012) who reported the significance of LRRXII family genes in providing resistance against necrotic fungi in tomato genotype. In hot pepper three genes (CaRLK23, 24 and 25) from subfamily LRK10L2, two genes (CaRLK15, 16) from SD1a subfamily and CaRLK 25, 26 from WAKLRK10L1 had showed significant up-regulation in resistant variety particularly during hyphae colonization and acervuli formation. Presence of *cis*-regulatory elements like TGACG (MeJA responsive), ERE (Ethylene stress responses) STRE, MYB and MYC (defense responsive) in promoter region are majorly involved in JA-ET pathway may be responsible for providing resistance. Chowdhury et al. (2017) [41]. Reported an increase in Jasmonic Acid (JA) and Ethylene (ET) hormone-mediated signaling pathways during biotrophic and necrotrophic phase of *Colletotrichum* infection which was responsible for governing disease resistance in sesame. Three genes CaRLK 17, 18, 19 from SD3 subfamily with *cis*-regulatory element like TCA element involved in salicylic acid regulation had showed significant up-regulation only during hyphae colonization. Qi et al. (2012) [42] reported the role of SA in hyphae growth and basal defense response. While Genes from SD2b family didn’t exhibit any up-regulation. Few genes CaRLK 2, 7, 22 and 17 from various subfamilies had showed significant downregulation. In rice among four WAK members, three genes OsWAK14, OsWAK91 and OsWAK92 act as positive regulators and OsWAK112d as negative regulator while providing quantitative resistance against *magnaporthae oryzae* [43]. Expression studies by Rt-qPCR were used to identify candidate genes / functional markers majorly involved in crop improvement programs.

**Fig. 4.**
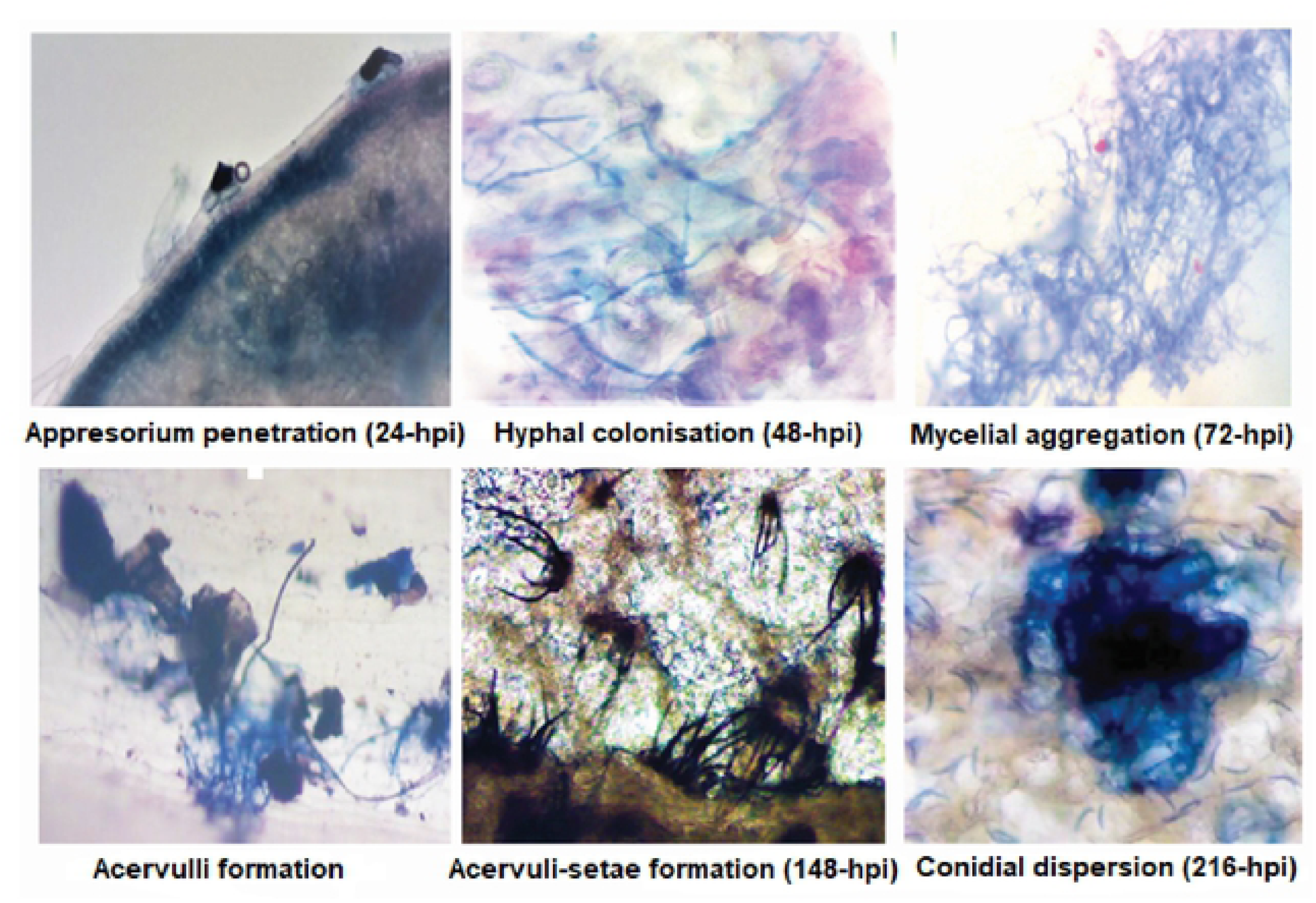
Histochemical observation of *Colletotrichum truncatum* infective structures on hot pepper leaves under electron microscope at 40 X magnification.

## Conclusion

This is the first report of genome-wide identification, characterization, and expression profiling of the non-RD kinase gene family in hot pepper. We had systematically analyzed and identified 26 CaRLKs, and characterized those using bioinformatics and expression analyses in response to *Colletotrichum truncatum* stress conditions. After nine days of infection one gene (CaRLK1) from LRRXII subfamily out of 26 non-RD genes was found to be expressed more in resistant genotype (PBC-80) than in susceptible genotype (Pusajwala). Moreover, identification of *cis*-elements in this gene enabled us to understand their role in conferring resistance in PBC-80 genotype by activation of the phytohormone JA-ET signaling pathway. Therefore this comprehensive analysis serves as a central platform to understand various physiological and biochemical functions performed by CaRLK1 gene in providing resistance. This investigation can also pave a way to analyze stress responsive gene expression studies in various pepper tissues and aids in promoting sustainable agriculture by using different crop improvement methods

## Acknowledgment

Authors are thankful to departmental heads and Co-scholars of bioinformatics lab for providing facilities to carry out the work.

## Supplementary data

All data associated with this paper can be found within the supplementary files.

## Supporting information

S1 Table. List of primers used for RLK expression analysis

S2 Table. Conserved motif based identification of Non RD kinases

S3 Table. Physiochemical properties of non-RD kinases in Hot pepper

S4 Table. Estimated Ka/Ks ratios and divergence times of the duplicated CaRLK genes

S5 Fig. Multiple sequence alignment of conserved motifs in kinase region of Hot pepper S6 Fig. Identified conserved motifs in non-RD RLK in hot pepper.

S7 Fig. Distribution of non-RD kinases on hot pepper chromosomes. Chromosome numbers are indicated at top of each bar. Scale is represented in mega bases (Mb).

S8 Fig. Exon-intron structure of non-RD kinases. Yellow indicates exon, black line indicates intron.

S9 Fig. *Cis-*acting elements in the promoter regions of CaRLK genes. S10 Fig. Sequence length cutoffs to build the credible set

